# Different vaccine platforms result in distinct antibody responses to the same antigen in haemodialysis patients

**DOI:** 10.1101/2024.01.14.575569

**Authors:** Nadya Wall, Rachel Lamerton, Fiona Ashford, Marisol Perez-Toledo, Aleksandra Jasiulewicz, Gemma D Banham, Maddy L Newby, Sian Faustini, Alex Richter, Haresh Selvaskandan, Roseanne E Billany, Sherna Adenwalla, Ian R Henderson, Max Crispin, Matthew Graham-Brown, Lorraine Harper, Adam F Cunningham

**Affiliations:** Institute of Applied Health Research, College of Medical and Dental Sciences, University of Birmingham, UK; Institute of Immunology and Immunotherapy, College of Medical and Dental Sciences, University of Birmingham, UK; University Hospitals Birmingham NHS Foundation Trust, Birmingham, UK; University Hospitals Coventry and Warwickshire NHS Trust, Coventry, UK; Department of Cardiovascular Sciences, University of Leicester, Leicester, UK; National Institute for Health Research (NIHR) Leicester Biomedical Research Centre, University Hospitals of Leicester NHS Trust and University of Leicester, Leicester, UK; School of Biological Sciences, University of Southampton, Southampton, UK; Institute for Molecular Bioscience, The University of Queensland, Brisbane, Australia

**Keywords:** Haemodialysis, Antibody, Immune Responses, SARS-CoV-2

## Abstract

Generalised immune dysfunction in chronic kidney disease, especially in patients requiring haemodialysis (HD), significantly enhances the risk of severe infections. Moreover, vaccine-induced immunity is typically reduced in HD populations, but the full mechanisms behind this remain unclear. The SARS-CoV-2 pandemic provided an opportunity to examine the magnitude and functionality of antibody responses in HD patients to a previously unencountered antigen, Spike (S)-glycoprotein, after vaccination with different vaccine platforms (viral vector (VV); mRNA (mRV)). Here, we compared total and functional anti-S antibody responses (cross-variant neutralisation and complement binding) in 187 HD patients and 43 healthy controls 21-28 days after serial immunisation. After 2 doses of the same vaccine, HD patients had anti-S antibody levels and complement binding capacity comparable to controls. However, 2 doses of mRV induced greater polyfunctional antibody responses than VV, yet previous SARS-CoV-2 infection or an mRV boost after 2 doses of VV significantly enhanced antibody functionality in HD patients. Therefore, HD patients can generate near-normal, functional antigen-specific antibody responses following serial vaccination to a novel antigen, suggesting largely intact B cell memory. Encouragingly, exploiting immunological memory by using mRNA vaccines and boosting may improve the success of vaccination strategies in this vulnerable patient population.

## Introduction

Patients with chronic kidney disease (CKD), particularly those requiring haemodialysis (HD), have a significantly greater risk of infection and poorer infection-related outcomes than the general population [1–4]. The nature and cause(s) of the secondary immunodeficiency state associated with CKD remain incompletely understood [5]. Although defects in innate and adaptive immunity have been described [6, 7], their contributions to the increased infection susceptibility in CKD/HD have not been fully characterised. Additionally, CKD patients have antibody responses to antigens such as tetanus toxoid, *Salmonella enterica* or cytomegalovirus that are comparable to non-CKD populations [8]. Since these antigens are likely first encountered relatively early in life, it indicates that CKD patients can maintain long-lived plasma cell responses against multiple antigens. This suggests that CKD/HD patients have dysfunctional immunity rather than being profoundly immunodeficient.

Vaccination is one of the few intervention available to modify the risk of infection and associated morbidity/mortality across populations, particularly for those with CKD/HD. Multiple studies show impaired seroconversion following various conventional vaccines in CKD and HD populations [9, 10], but this is not universal [11, 12]. Respiratory infections constitute a high proportion of the infection burden in CKD patients [2, 13], but evaluating immune responses to seasonal vaccines e.g. influenza/pneumococcus is complex due to unquantifiable previous exposures to these common pathogens. Studies often use serological responses as a surrogate for vaccine effectiveness in reducing infection risk/severity on a population level, but antibody correlates of protection on an individual level are less clear.

During the SARS-CoV-2 pandemic the global population was exposed to a novel pathogen. The pandemic tragically highlighted the markedly increased burden of infection and associated mortality in patients with CKD, particularly in those requiring HD. In HD patients, symptomatic presentations were frequently associated with poor outcomes (25-30% case fatality rate in pre-vaccination waves [14, 15]). However, subsequent sero-epidemiological studies unexpectedly found that some HD patients had encountered this novel pathogen and were able to generate SARS-CoV-2-specific immune responses without experiencing symptoms [16]. This points to the potential conservation, in a proportion of HD patients, of the capacity to induce *de novo* protective immunity, even to pathogens capable of causing severe disease.

Clinical outcomes and infection susceptibility in HD patients were significantly improved with the roll-out of SARS-CoV-2 vaccines [17–22], albeit vaccine efficacy is generally reported to be lower than in the general population [23, 24]. This emphasises the real possibility of positively modulating immune responses in HD patients to improve infection-related outcomes. Fundamental to this is the need for better understanding of where immune deficiencies lie in these patients and how they result in dysregulated control of pathogens and infections.

Immune responses to spike glycoprotein (S) alone can be sufficient to protect against SARS-CoV-2 infection, hence all licensed vaccines against SARS-CoV-2 depend upon delivery of S to the immune system [25]. Nevertheless, the mechanism through which this antigen is delivered to the immune system differs. The most widely used vaccine platforms to deliver S are vector-based or mRNA technologies. These different vaccine platforms allow us to assess the nature of the immune response induced to a novel antigen and any potential influence of the vaccine platform and exposure to the pathogen itself on these responses. Collectively this can provide insights into what protective immunity looks like within the HD patient population and guide how best to manipulate their immune system to reduce their risk of infection and improve infection-related outcomes.

In this study, we systematically characterise humoral immunity in patients requiring HD by examining quantitative and qualitative antibody responses to SARS-CoV-2 S antigen after serial vaccination with 2 different vaccine platforms, in the context of previous SARS-CoV-2 infection history.

## Results

### Demographics/descriptors of study population

Demographics and clinical parameters of study participants are described in Table 1. Patients requiring HD were significantly older, more likely to be male and of non-white ethnicity than controls. As expected, HD patients had more comorbidities than controls, with a significantly greater prevalence of diabetes mellitus (DM, 41% versus nil in controls) and immunosuppression (IS, 9% in HD versus nil in controls). Patients requiring HD were broadly similar across the two UK study sites. Eighteen HD patients developed post-vaccination SARS-CoV-2 infection – representing 1 in 10 at both sites (Table 1).

**Table 1.**
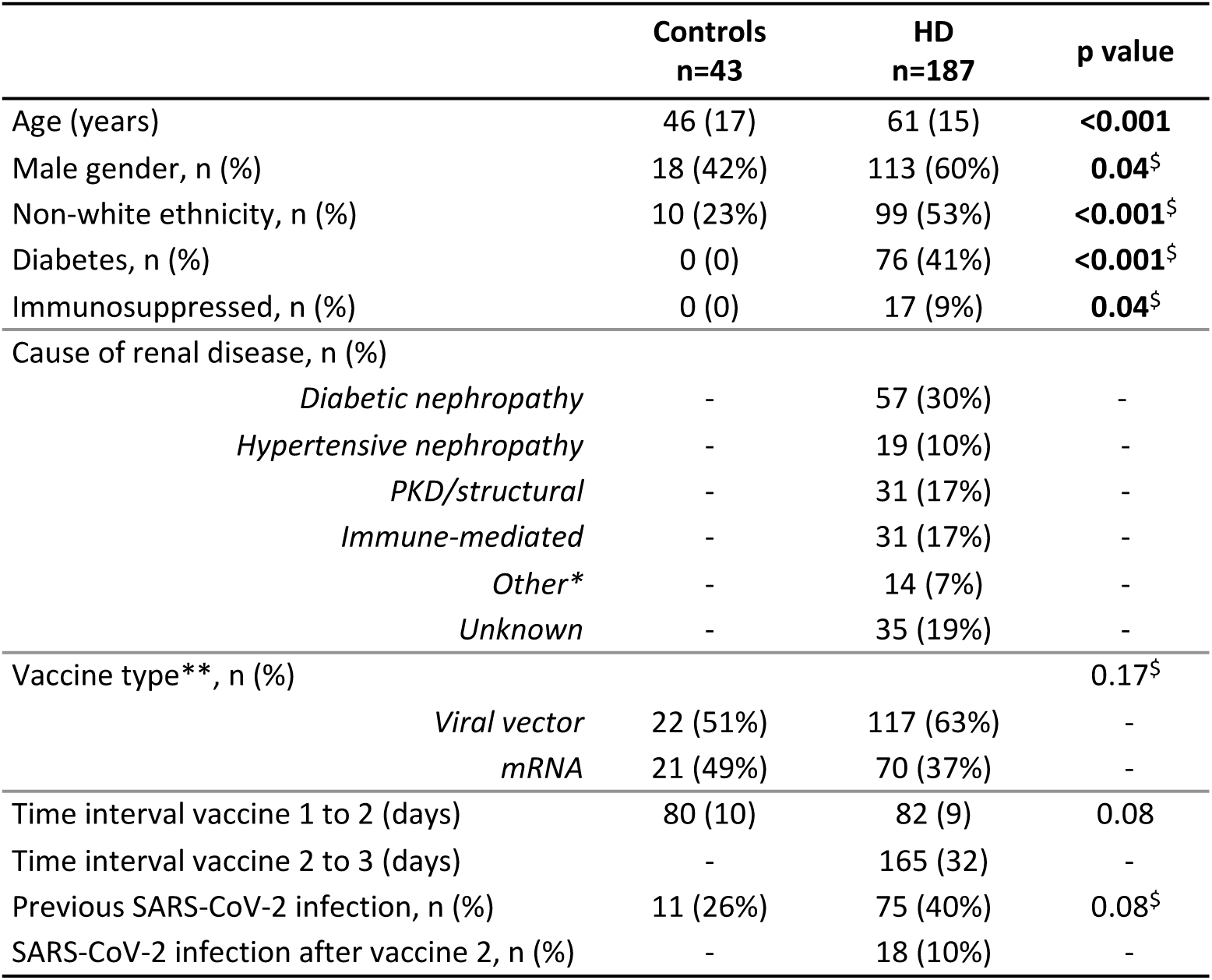
Demographics and clinical descriptors of study population. . Data presented as median (IQR), unless stated otherwise. Statistical comparisons performed using Mann Whitney U test for continuous data and Fisher’s exact test (denoted by ^$^) for categorical data. p value <0.05 considered significant (highlighted in bold typeface). *Other causes of renal disease include multiple myeloma, trauma and renal tuberculosis. **denotes vaccine type given for first 2 doses. *Abbreviations: HD – haemodialysis, PKD – polycystic kidney disease*.

A greater proportion of HD patients than controls received 2 doses of AZD1222 vaccine than BNT162b2 (hereafter described as viral vector (VV) and mRNA vaccines (mRV), respectively), but this did not reach statistical significance, and the time interval between first and second vaccine doses was similar between controls and HD patients. A higher proportion of HD patients had previous exposure to SARS-CoV-2 than controls (defined as either previous positive nasopharyngeal SARS-CoV-2 PCR and/or detection of anti-nucleocapsid (anti-N) antibody at study entry), but this was not statistically significant. As such, age, gender, ethnicity, HD status, vaccine type and previous SARS-CoV-2 exposure were included as co-variables in subsequent multivariable analyses. All HD patients received mRV as their third vaccine dose.

### Patients requiring HD generate similar quantitative antibody responses as controls following 2 vaccine doses

HD patients and controls had similar anti-S antibody levels at 21-28 days after a second vaccine dose, when stratified by vaccine type and prior SARS-CoV-2 infection (Figure 1 A-C, Table 2). Using the assay positive threshold of 0.8U/ml for anti-S levels, all controls and almost all HD patients (n=183/187, 98%) were seropositive after 2 vaccine doses.

**Figure 1.**
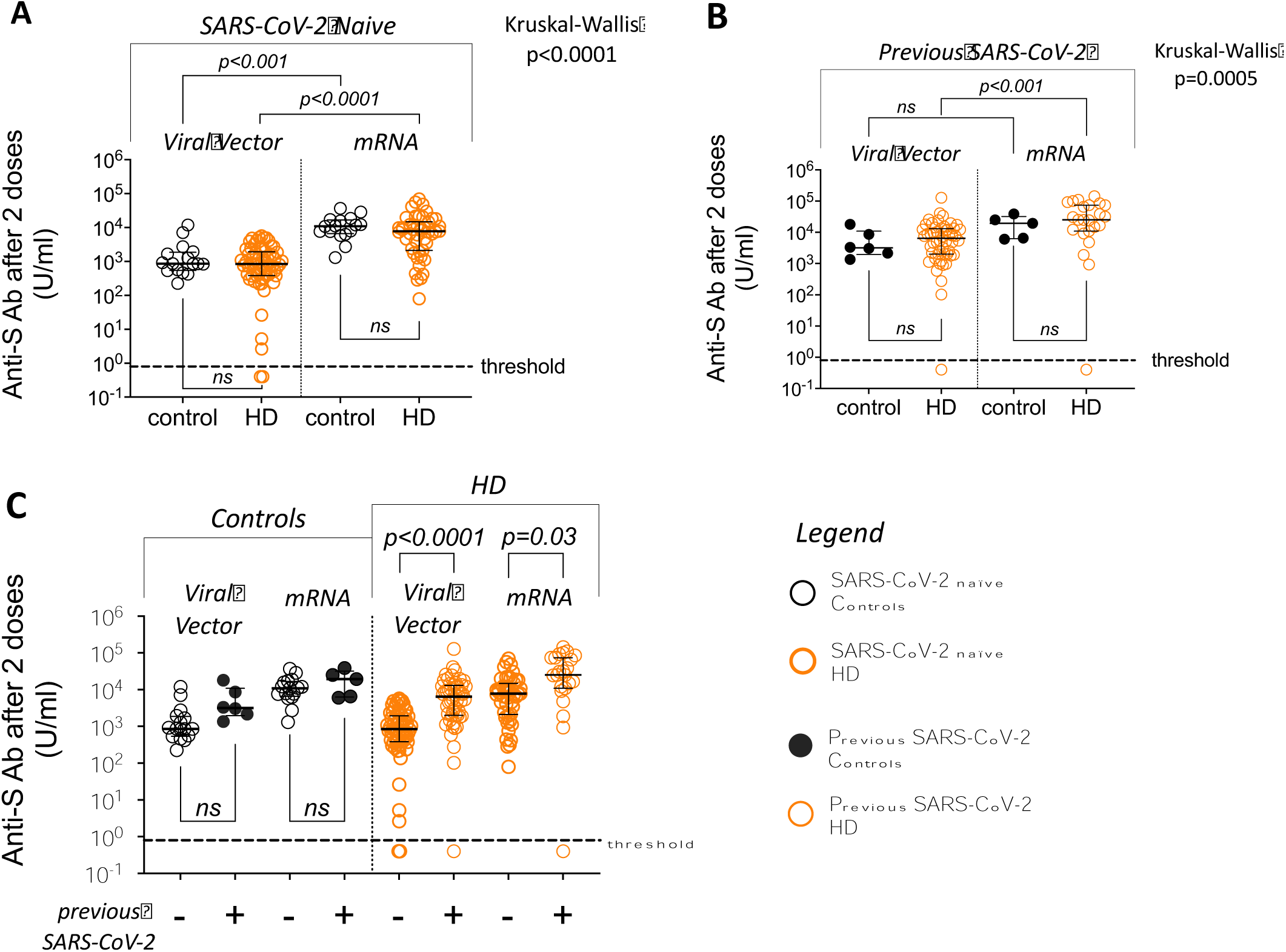
Antigen-specific antibody in sera collected 21-28 days after 2 doses of SARS-CoV-2 vaccine – comparison between controls and patients requiring HD. Comparisons of anti-S antibody levels between controls (black symbols) and patients requiring HD (orange symbols) – shown in legend. Filled symbols represent individuals with previous SARS-CoV-2 infection. Blue columns denote data from mRNA vaccinees. A – data from SARS-CoV-2 naïve individuals only; B – data from individuals with previous SARS-CoV-2 infection only; C – all data shown. Threshold level of antibody detection (“seropositivity”) shown as dashed line (0.8U/ml). Kruskal-Wallis test used to compare groups in panels A and B, with post-hoc Dunn’s multiple comparison p values shown (ns denotes not significant, p>0.05). Mann-Whitney U test used for pre-defined comparisons within groups in panel C.

**Table 2.**
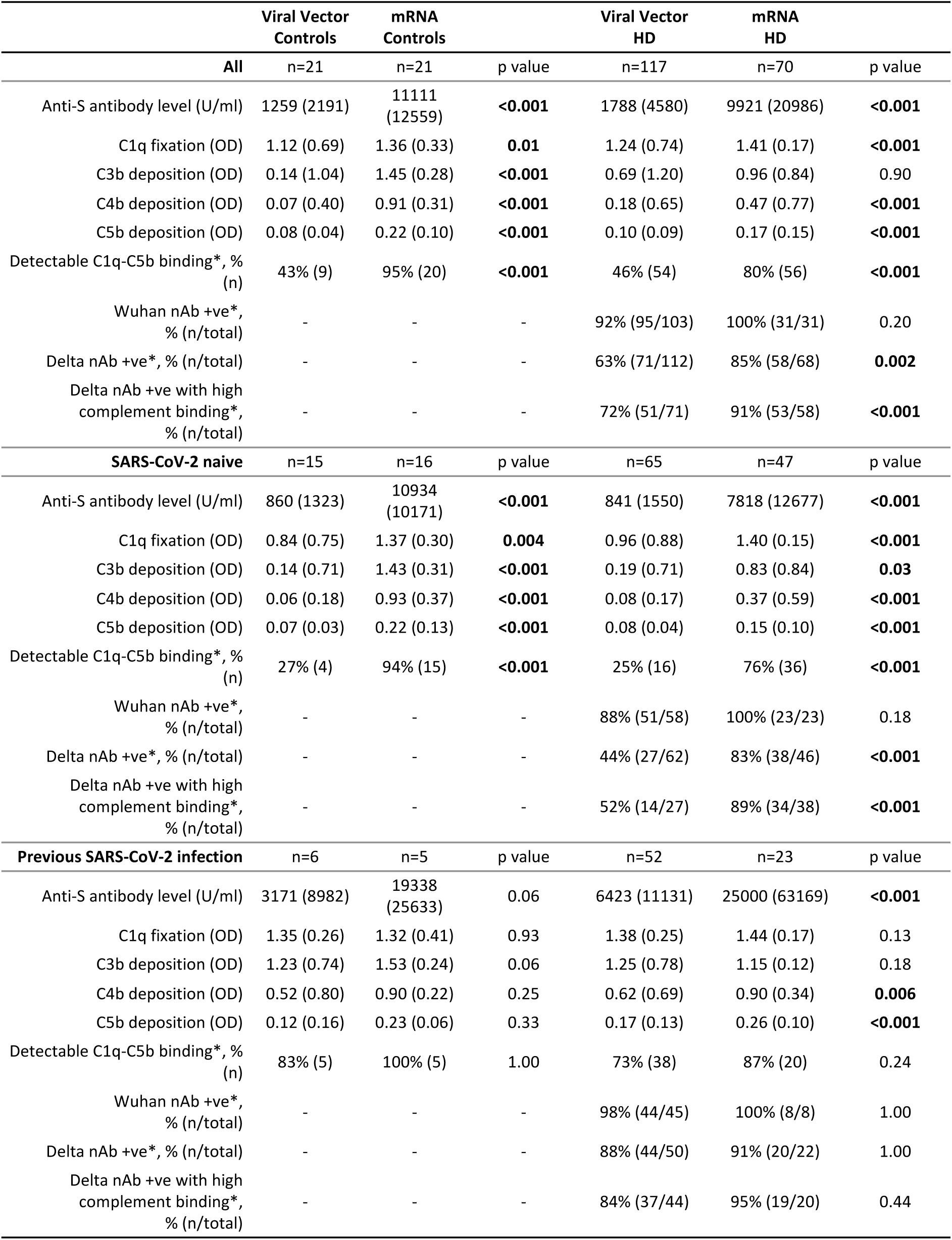
Serology after 2 vaccine doses - comparison between controls and HD patients. Initially, comparisons between the viral vector and mRNA vaccines were performed for the control and HD patients separately. The analyses were then repeated for the subgroups of patients that were SARS-CoV-2 naïve at the time of study entry, and those that had a previous SARS-CoV-2 infection. Data are presented as median (IQR) unless otherwise stated. For proportions - denominator (total n) is given where there is missing data. Mann-Whitney U test p values shown for continuous data, Fisher’s exact test p values shown for categorical data (denoted by *). P values <0.05 considered as significant and highlighted in bold typeface. *Abbreviations: nAb – neutralizing antibody activity (positivity defined as IC50 40 or greater)*.

Within both groups, post-vaccination anti-S antibody levels were generally higher in individuals vaccinated with 2 doses of mRV than VV, particularly in SARS-CoV-2 naïve individuals (Figure 1A, B). Previous infection with SARS-CoV-2 was associated with higher anti-S levels for HD patients receiving 2 vaccine doses, independent of vaccine type (Figure 1C). For analyses performed in HD patients, mRV, previous SARS-CoV-2 infection and non-immunosuppressed status were significant predictors of higher anti-S antibody levels in a multivariable linear regression model that also included age, gender, ethnicity and diabetes (Supplementary Table 1).

In summary, we show that, after 2 vaccine doses, HD patients can have similar antigen-specific antibody levels to novel antigen as controls matched for vaccine type and previous pathogen exposure.

### Vaccine platform and HD influence fixation and deposition of complement by antigen-specific antibody

As quantitative antigen-specific antibody responses after 2 vaccine doses were largely similar between controls and HD patients, we then compared surrogates of antigen-specific antibody functionality between the groups. One such surrogate is the binding and activation of complement components by antigen-specific antibody - fixation of C1q and deposition of downstream products of the complement cascade (C4b, C3b and C5b) [26].

In seropositive individuals, detection of complement component binding in our solid-phase assay positively correlated with the quantity of circulating anti-S antibody (Figure 2A-D). As such, both mRV and previous SARS-CoV-2 infection were significantly associated with greater binding of complement components by anti-S antibody (Table 2, Figure 3A-D).

**Figure 2.**
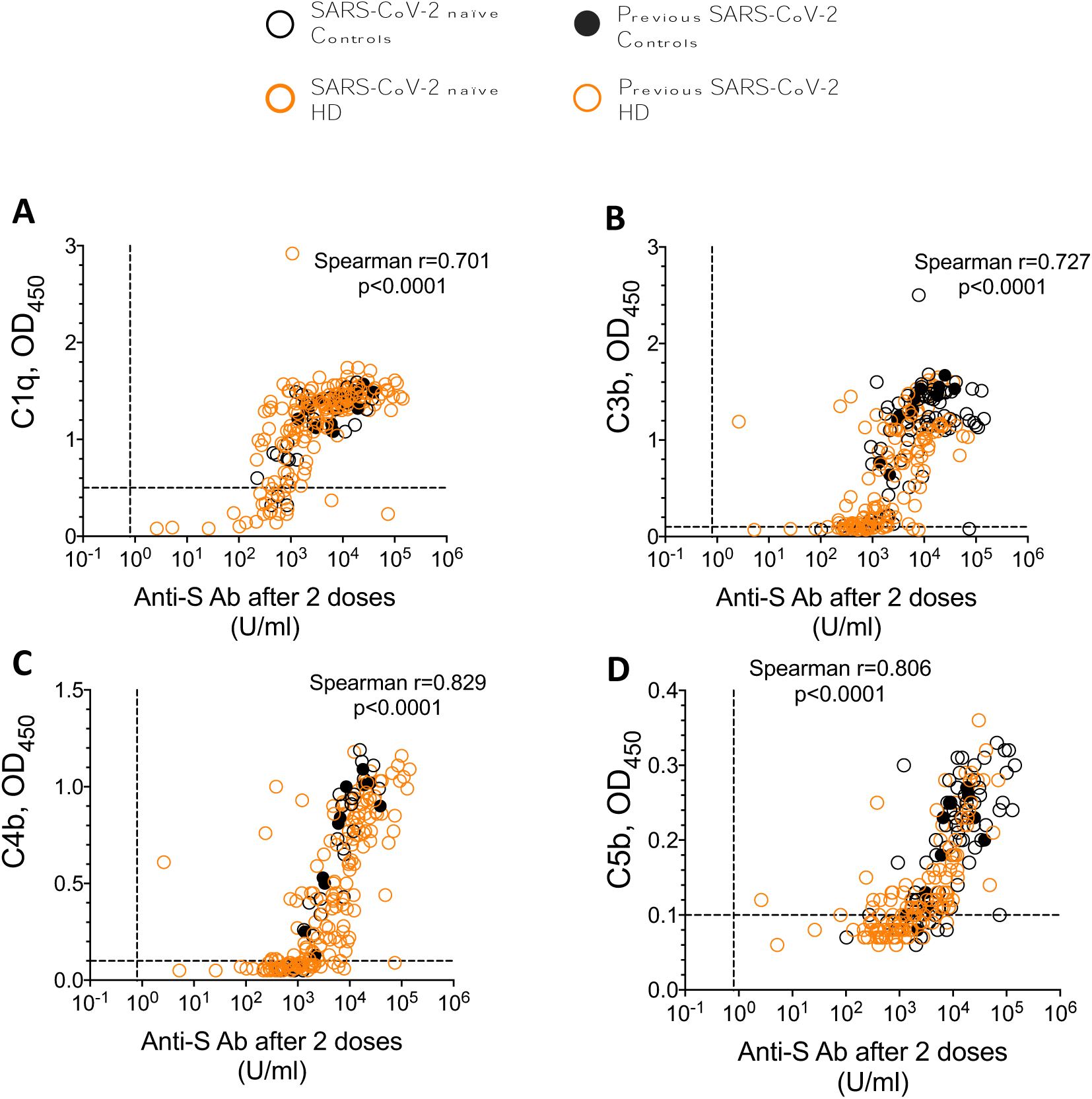
Correlation between anti-S antibody levels and binding of complement components in controls and patients requiring HD. Data shown for all seropositive individuals – **A-D** correspond to C1q, C3b, C4b and C5b, respectively. Spearman’s rank correlation coefficient and p value shown for all data points. Black symbols denote controls, orange symbols denote HD patients and filled symbols denote previous SARS-CoV-2 infection – see Legend. Threshold level of antibody detection (“seropositivity”) shown as vertical dashed line (0.8U/ml), complement binding assay negative thresholds shown as horizontal dashed lines (0.5 for C1q and 0.1 for C3b/4b/5b).

**Figure 3.**
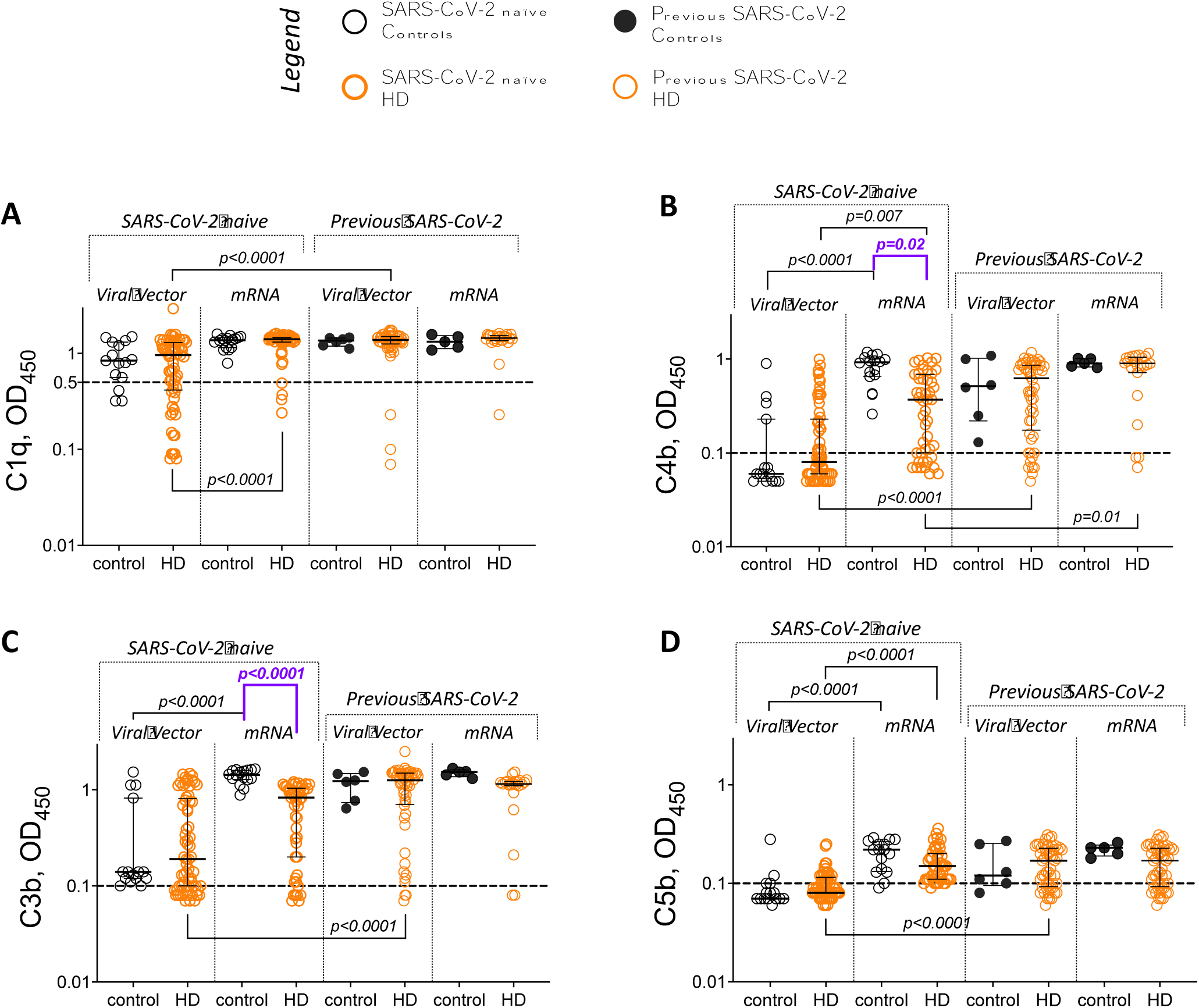
Complement component binding by antigen-specific antibody in sera collected 21-28 days after 2 doses of SARS-CoV-2 vaccine – comparison between controls and patients requiring HD. Comparisons shown between controls (black symbols) and patients requiring HD (orange symbols) with data grouped by vaccine type and previous SARS-CoV-2 exposure (filled symbols denote previous infection); **A-D** correspond to C1q, C3b, C4b and C5b, respectively. Assay negative threshold (0.1 for C3b-C5b, 0.5 for C1q) shown as dashed line. Kruskal-Wallis test used for comparisons between groups (p<0.05 for all complement components, not shown), with post-hoc Dunn’s multiple comparison p values shown; Purple colour in panels I and J denotes comparisons of particular interest (described in main text); ns denotes p>0.05.

Although the majority of seropositive individuals (controls and HD patients) had detectable C1q fixation, SARS-CoV-2 infection-naïve VV recipients generally showed lower binding of C3b-C5b by anti-S antibody than their mRV vaccinated counterparts (Table 2, Figure 3 B-D). Indeed, a significantly lower proportion of SARS-CoV-2 infection-naïve VV recipients had detectable binding of all 4 complement components tested, both in control and HD cohorts (Table 2, Fisher’s exact p<0.001).

Overall, when stratified by vaccine type and previous SARS-CoV-2 exposure, patients requiring HD exhibited greater heterogeneity than controls in the levels of complement binding detected (Figure 3A-D). Interestingly, despite having similar anti-S levels, SARS-CoV-2 naïve mRV vaccinated HD patients showed significantly lower deposition of C4b and C3b than controls matched for vaccine type and previous pathogen exposure (Figure 3 B,C – comparisons highlighted in purple).

In summary, we show that complement component fixation and deposition by antigen-specific antibody is highest in individuals that received 2 doses of mRV or those that have had a previous infection, with HD patients showing greater variability in responses than controls.

### In HD patients, the mRNA vaccine induces a broader functionality of antigen-specific antibodies than the viral vector vaccine

Sera from HD patients can exhibit reduced neutralisation activity against SARS-CoV-2 when compared to healthy controls [27]. Neutralisation activity against related, but genetically distinct SARS-CoV-2 variants reflects the breadth of immune responses induced and is a desirable feature of the antibody response to SARS-CoV-2 vaccination [28–30]. As such, we then assessed neutralising activity of sera (nAb) against the Delta variant as a proxy of Fab segment diversity and functionality.

Most patients requiring HD for whom data was available were able to neutralise the Wuhan SARS-CoV-2 variant after 2 doses of any SARS-CoV-2 vaccine (94%, n=126 of 134, Table 2). For the Delta variant, the mRV platform was associated with a greater prevalence of detectable nAb than the VV vaccine in SARS-CoV-2 infection-naïve HD patients (Table 2, Supplementary Figure 1A, Fisher’s exact 2-tailed p<0.0001), but no differences by vaccine type were observed in those who had experienced SARS-CoV-2 infection. Delta nAb positive HD patients had significantly higher anti-S levels than Delta nAb negative patients (Supplementary Figure 1B).

We then cross-compared Delta variant neutralisation and binding of complement components as surrogate measures of Fab and Fc segment antibody function, respectively. Surprisingly, detectable Delta nAb and complement component binding were not mutually inclusive in HD patients. This was most pronounced in SARS-CoV-2-naïve VV vaccinees, where almost half of individuals with detectable Delta neutralising antibody did not have detectable binding of all 4 complement components tested (Table 2). This proportion was significantly lower than that seen in SARS-CoV-2 naïve mRV and VV vaccinees with previous SARS-CoV-2 infection (Table 2, Fisher’s exact 2-tailed p<0.001 and p=0.02, respectively).

Interestingly, the presence of highly functional anti-S antibody (either detectable Delta nAb and/or binding to all 4 complement components tested) was significantly associated with reduced likelihood of post-vaccination SARS-CoV-2 infection, independent of age, gender, ethnicity, presence of DM/immunosuppression, vaccine type and HD centre (included due to different infection screening practices) (Supplementary Table 2).

Overall, vaccination with the mRV platform in HD patients is associated with greater antibody functionality than with VV and the presence of highly functional antigen-specific antibody is associated with protection against post-vaccination infection.

### mRNA vaccination after 2 doses of a viral vector vaccine significantly improves antigen-specific antibody function in patients requiring HD

During the study period, the HD population (as a clinically vulnerable group) were offered a third vaccine dose (booster) at 6 months after completion of the primary (2 dose) course, but this was not routinely done for controls [31]. This enabled us to compare the impact of receiving heterologous (VV, followed by mRV) or homologous (mRV only) vaccines on quantitative and functional measures of circulating anti-S antibody in patients requiring HD.

All HD patients had detectable anti-S antibody after 3 doses of SARS-CoV-2 (Figure 4). The third vaccine dose significantly increased anti-S antibody levels in the viral vector vaccine groups, irrespective of previous SARS-CoV-2 infection history (Figure 4, Table 3). In HD patients who had previously received mRV, only the group without previous SARS-CoV-2 infection demonstrated an increase in antibody levels after the third vaccine dose (Figure 4, Table 3).

**Figure 4.**
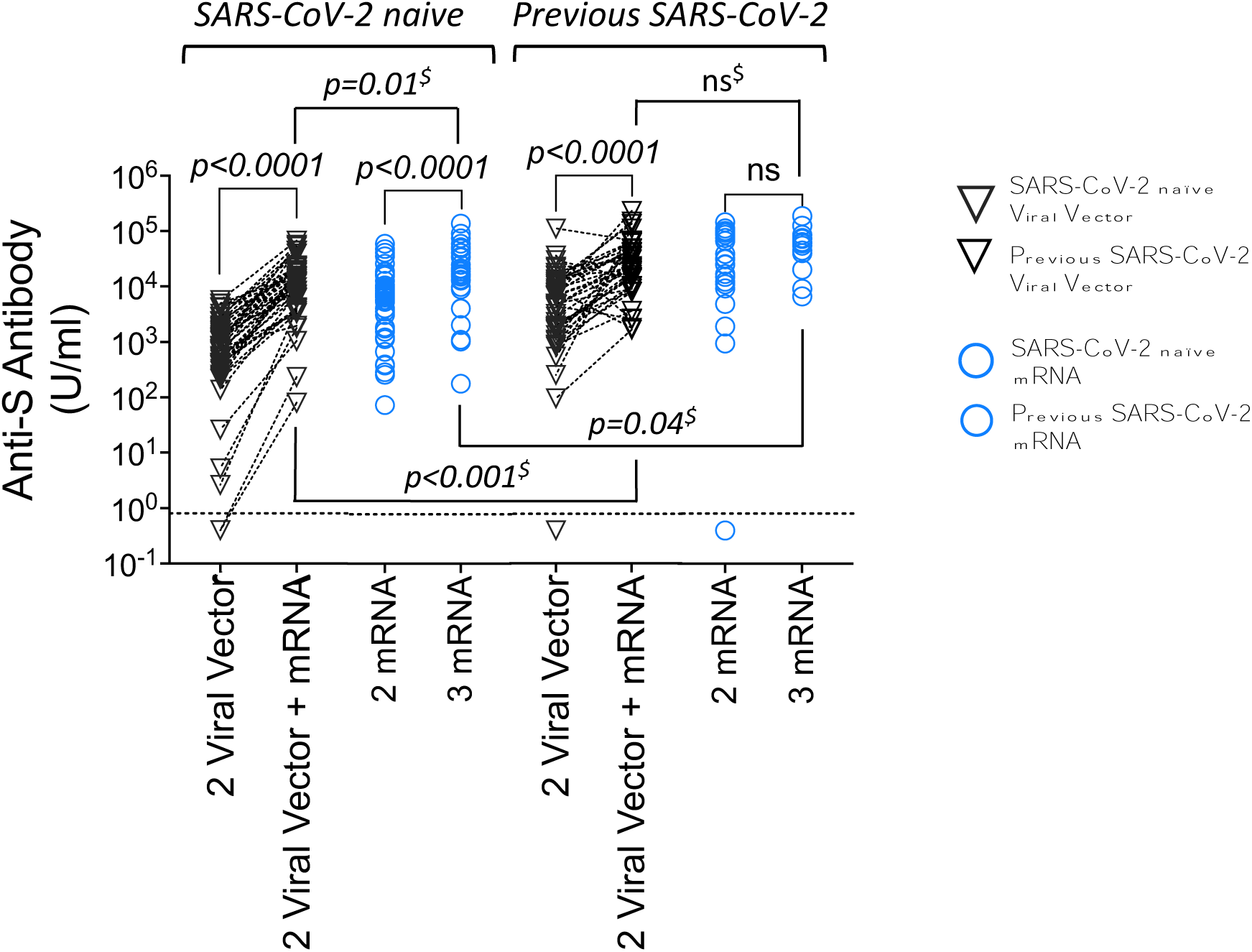
– Anti-S antibody levels in sera of HD patients 21-28 days after 2 and 3 vaccine doses. Anti-S antibody levels in HD patients after 2 and 3 vaccine doses – comparisons by first vaccine type (grey – viral vector, blue – mRNA), split by previous SARS-CoV-2 infection (filled symbols denote previous infection) – see Legend. Paired statistical comparisons performed between timepoints using Wilcoxon signed-rank test (post-dose 2 v post-dose 3 levels) and Mann Whitney test U used to compare antibody levels at the same time points by previous SARS-CoV-2 exposure (denoted by ^$^).

**Table 3.** – Comparison of serology parameters after 2 and 3 vaccine doses in patients requiring HD, stratified by previous SARS-CoV-2 exposure. Initially, comparisons between 2 and 3 vaccine doses were performed separately for HD patients initially vaccinated with viral vector and mRNA vaccines. The analyses were then repeated for the subgroups of patients that were SARS-CoV-2 naïve at the time of the first vaccine, and those that had a previous SARS-CoV-2 infection. Data presented as median (IQR) unless otherwise stated. For proportions - denominator (total n) given where there is missing data. Mann-Whitney U test p values shown for continuous data, Fisher’s exact test p values shown for categorical data (denoted by *). P values <0.05 considered as significant and highlighted in bold typeface. *Abbreviations: nAb – neutralizing antibody activity (positivity defined as IC50 40 or greater)*.

|  | Viral Vector |  |  | mRNA |  |  |
| --- | --- | --- | --- | --- | --- | --- |
| All HD patients | 2 doses<br>n=117 | 2 doses+mRNA<br>n=92 | p value | 2 doses<br>n=70 | 2 doses+mRNA<br>n=40 | p value |
| anti-S Ab level (IU) | 1788 (4580) | 18227 (36914) | **<0.001** | 9921 (20986) | 26967 (45573) | **<0.001** |
| fold change in anti-S Ab level | ref | 9.9 (22.8) |  | ref | 3.3 (6.3) |  |
| C1q fixation (OD) | 1.23 (0.74) | 1.41 (0.28) | **<0.001** | 1.41 (0.17) | 1.40 (0.30) | 0.90 |
| C3b deposition (OD) | 0.69 (1.20) | 1.29 (0.24) | **<0.001** | 0.96 (0.84) | 1.22 (0.54) | **<0.001** |
| C4b deposition (OD) | 0.18 (0.65) | 0.90 (0.42) | **<0.001** | 0.47 (0.77) | 0.74 (0.66) | **<0.001** |
| C5b deposition (OD) | 0.10 (0.09) | 0.31 (0.13) | **<0.001** | 0.17 (0.15) | 0.24 (0.15) | **<0.001** |
| Detectable C1q-C5b binding*, % (n) | 46% (54) | 95% (87) | **<0.0001** | 80% (56) | 85% (34) | 0.61 |
| Delta nAb +ve*, % (n/total) | 63%<br>(71/112) | 98% (83/85) | **<0.0001** | 85% (58/68) | 100% (23/23) | 0.06 |
| Delta nAb +ve with high complement binding*, % (n/total Delta nAb +ve) | 72%<br>(51/71) | 98% (81/83) | **<0.0001** | 91% (53/58) | 83% (19/23) | 0.26 |
| SARS-CoV-2 naïve | 2 doses<br>n=65 | 2 doses+mRNA<br>n=46 | p value | 2 doses<br>n=47 | 2 doses+mRNA<br>n=28 | p value |
| anti-S Ab level (IU) | 841 (1550) | 12401 (17966) | **<0.001** | 7818 (12677) | 20800 (35949) | **<0.001** |
| fold change in anti-S Ab level | ref | 14.6 (33.1) |  | ref | 3.7 (6.4) |  |
| C1q fixation (OD) | 0.96 (0.88) | 1.37 (0.34) | **<0.001** | 1.40 (0.15) | 1.32 (0.34) | 0.95 |
| C3b deposition (OD) | 0.19 (0.71) | 1.16 (0.34) | **<0.001** | 0.83 (0.84) | 1.09 (0.76) | **<0.001** |
| C4b deposition (OD) | 0.08 (0.17) | 0.67 (0.55) | **<0.001** | 0.37 (0.59) | 0.46 (0.71) | **0.002** |
| C5b deposition (OD) | 0.08 (0.04) | 0.26 (0.17) | **<0.001** | 0.15 (0.10) | 0.17 (0.14) | **0.002** |
| Detectable C1q-C5b binding*, % (n) | 25% (16) | 89% (41) | **<0.0001** | 77% (36) | 82% (23) | 0.77 |
| Delta nAb +ve*, % (n/total) | 44%<br>(27/62) | 95% (40/42) | **<0.0001** | 83% (38/46) | 100% (19/19) | 0.09 |
| Delta nAb +ve with high complement binding*, % (n/total Delta nAb +ve) | 52%<br>(14/27) | 95% (38/40) | **<0.0001** | 89% (34/38) | 79% (15/19) | 0.42 |
| Previous SARS-CoV-2 infection | 2 doses<br>n=52 | 2 doses+mRNA<br>n=46 | p value | 2 doses<br>n=23 | 2 doses+mRNA<br>n=12 | p value |
| anti-S Ab level (IU) | 6423<br>(11131) | 30650 (46987) | **<0.001** | 25000 (63169) | 56589 (57269) | 0.11 |
| fold change in anti-S Ab level | ref | 6.8 (9.9) |  | ref | 2.2 (4.0) |  |
| C1q fixation (OD) | 1.38 (0.25) | 1.45 (0.60) | **<0.001** | 1.44 (0.17) | 1.48 (0.26) | 0.70 |
| C3b deposition (OD) | 1.25 (0.78) | 1.33 (0.16) | **0.009** | 1.15 (0.12) | 1.36 (0.20) | **0.03** |
| C4b deposition (OD) | 0.62 (0.69) | 0.99 (0.21) | **<0.001** | 0.90 (0.34) | 0.93 (0.24) | 0.21 |
| C5b deposition (OD) | 0.17 (0.13) | 0.34 (0.06) | **<0.001** | 0.26 (0.10) | 0.28 (0.07) | 0.10 |
| Detectable C1q-C5b binding*, % (n) | 73% (38) | 100% (46) | **<0.0001** | 87% (20) | 92% (11) | 0.99 |
| Delta nAb +ve*, % (n/total) | 88%<br>(44/50) | 100% (42/42) | **0.03** | 91% (20/22) | 100% (4/4) | 0.99 |
| Delta nAb +ve with high complement binding*, % (n/total Delta nAb +ve) | 84%<br>(37/44) | 100% (42/42) | **0.01** | 95% (19/20) | 100% (4/4) | 0.11 |

A third vaccine dose significantly increased the proportion of Delta nAb positive individuals in VV recipients (Table 3), such that, after 3 vaccine doses, there were no longer any vaccine type-associated differences in the capacity of HD patient sera to neutralise the Delta SARS-CoV-2 variant (Fisher’s exact p>0.99). A third vaccine dose also increased complement component binding in the HD patient cohort. This was most pronounced in VV vaccinees, regardless of previous SARS-CoV-2 exposure, where binding of C1q, C3b, C4b and C5b were significantly higher after 3 doses of vaccine than after 2 (Table 3).

A third vaccine dose had a striking impact on the antibody functionality of VV vaccinees as measured by dual capacity to neutralise the Delta variant and bind complement components (Figure 5A), particularly in SARS-CoV-2-naïve HD patients (Figure 5B). Here, the third vaccine increased the proportion of Delta neutralising, complement fixing antibody from just over 50% to 95%, a level similar to that seen in those that received triple mRV (Table 3).

**Figure 5.**
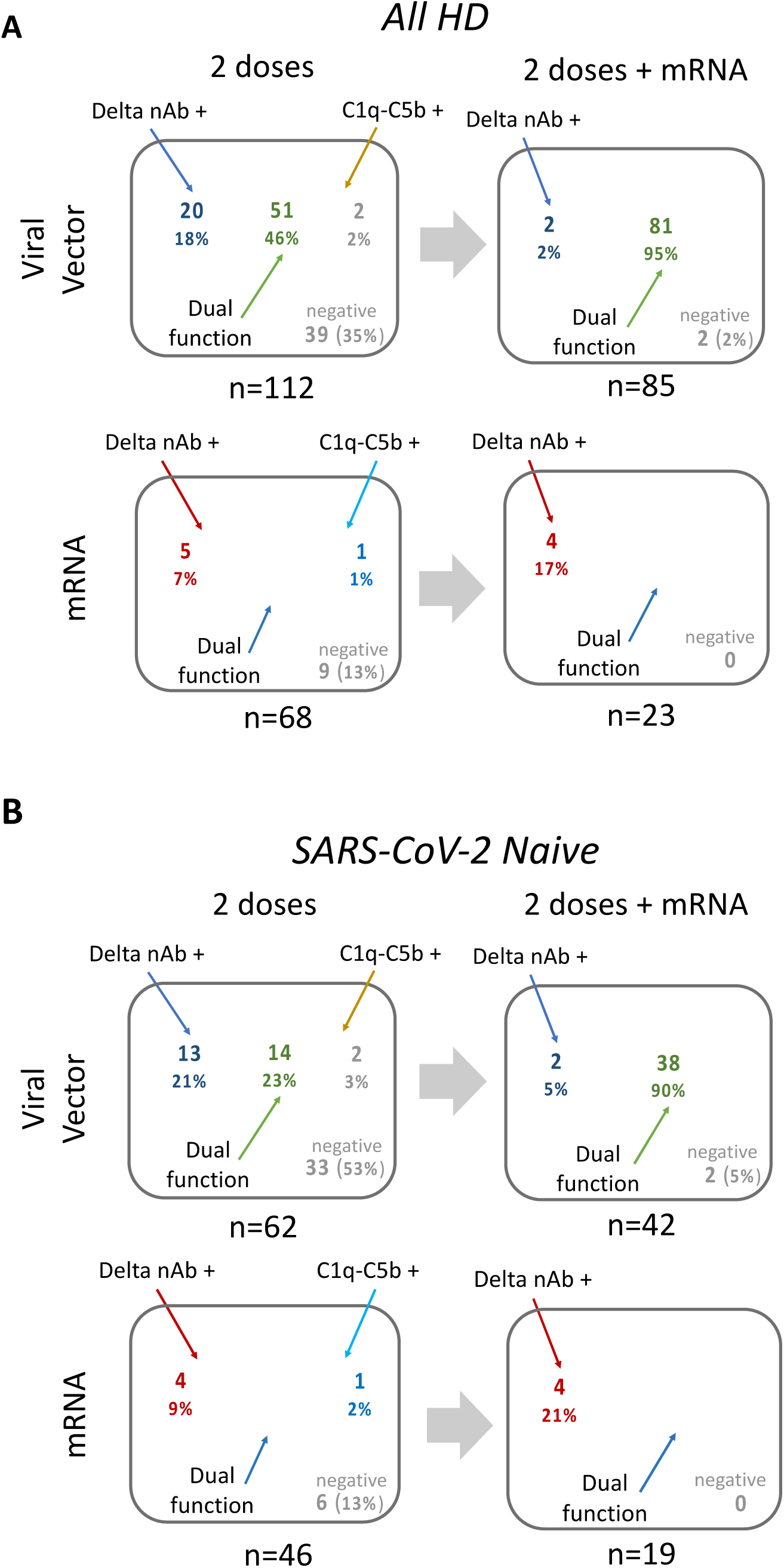
Antigen-specific antibody functionality in patients requiring HD – comparison between 2 and 3 vaccine doses. **A**: Diagrammatic representation of antigen-specific antibody functionality after 2 and 3 vaccine doses for all HD patients - comparisons by vaccine type. Venn diagrams showing overlap of neutralization activity against Delta VoC (Delta nAb +) with binding of all 4 complement components tested (C1q-C5b +) to denote antibody with dual function; n and % of total for whom data was available are shown. **B**: as above, but for SARS-CoV-2 naïve HD patients only.

In summary, an mRNA vaccine given after 2 doses of a VV vaccine significantly improves the quality of antigen-specific antibody in HD patients to levels comparable to individuals receiving mRNA vaccines alone.

## Discussion

In this study, we have examined the quantity and features of the quality of antigen-specific antibody following serial vaccination in patients requiring HD, a patient group with a secondary immunodeficiency and at high risk of infection. SARS-CoV-2 and its global spread, alongside the rapid development of vaccines, represented a unique opportunity to examine how immune responses to novel antigens develop in this patient population. Moreover, the development of different vaccine platforms allows us the opportunity to understand how immunity develops to the same antigen presented in different immunological contexts.

Patients requiring HD can generate anti-S antibody after SARS-CoV-2 infection, which confers protection from subsequent infection in survivors [32]. Nevertheless, after primary vaccination, quantitative humoral anti-S responses are typically lower than in controls [33–36]. This may be because HD patients, who often have a B cell lymphopenia, have a less diverse naïve B cell repertoire [37]. This could result in fewer B cells recruited into the primary response and yield stunted antibody responses, similar to what is observed in patients treated with B-cell depletion therapies [38]. However, our data show that subsequent recall humoral responses are largely preserved in HD patients, compared to equivalent controls, similar to that observed by others [19, 39–41]. This suggests that, although primary B cell responses may be different in HD patients, germinal centres can form and produce memory B cells that can respond upon subsequent antigen challenge, both in the context of vaccination and infection. As such, this suggests that HD patients have a largely intact immunological apparatus that can ultimately enable near-normal antibody responses, but this requires induction of immunological memory recall, combined with the optimised methods of antigen delivery (discussed further below).

Although the quantity of antigen-specific antibody is frequently used as the primary correlate of vaccine-associated immune responses and protection, antibody levels on their own do not always predict disease susceptibility [42]. As such, measures of antibody functionality offer insights into the quality or “success” of adaptive immune responses.

Antibodies consist of two functional domains - the highly variable antigen-binding fragment (Fab) conferring antigen specificity, together with the crystallizable fragment (Fc) that initiates downstream effector functions through interaction with complement proteins and Fc receptors (FcR). Virus binding and neutralisation represents a measure of Fab function. However, antibody neutralising activity alone may not always be enough to protect against disease [43, 44] and it may need to be enhanced by the activation of effector functions [45, 46]. Thus, the engagement of the classical complement pathway [47] may contribute to this or be a proxy for the development of antibodies with a broader functionality.

Little is understood about Fc function of antigen-specific antibody in patients requiring haemodialysis. Although we only assessed complement binding to antibody directed against Wuhan spike glycoprotein, others have shown Fc function to be conserved for anti-S antibody with cross-reactivity against different virus variants [48, 49]. The capacity of a vaccine to induce anti-S antibody capable of C1q fixation and downstream binding of C3b/4b/5b in HD patients shows that this functional property can be induced, but that the degree of binding is influenced by previous SARS-CoV-2 infection history and vaccine type.

SARS-CoV-2-naïve HD patients generate significantly lower nAb titres to the reference (Wuhan) virus and other variants than controls after 2 doses of a viral vector, but this difference is not observed for mRNA vaccinated groups [27]. In this study, we find a significant effect of previous SARS-CoV-2 infection in promoting the production of cross-variant neutralising antibody in HD patients, as has been shown for healthy individuals [50]. Combined with the other effects of prior infection on the antibody responses that we observed and the demonstrable impact of vaccine platform on these responses, it emphasises that antibody responses in HD patients can be “normal” or “near-normal” when the antigen is encountered in certain immunological contexts. The importance of this is that it indicates that vaccines can be tailored, either in how they are built and/or delivered, to improve the immune responses they provoke and, potentially, the clinical benefit they confer. This is emphasised by our finding that breadth of neutralisation (Fab function and potentially diversity) and complement binding (Fc function) are not mutually inclusive in patients requiring HD, but are influenced by the vaccine platform used. Boosting with a third mRV dose resulted in equivalent humoral responses in HD patients, independent of the vaccine type received previously, similar to what has been observed by others [51]. Again, this emphasises the importance of the maturation of antibody responses, presumably through further germinal centre induction, but may also indicate either that heterologous vaccination protocols can be beneficial in their own right or that mRNA vaccines offer some superiority in the HD population [24]. Some evidence for these points has already been noted in healthy populations [52], but further studies are needed to test these potential options in patients requiring HD.

Despite our HD cohort being relatively large, ethnically diverse and largely representative of the wider HD patient population in terms of comorbidity, our study has limitations. Our smaller control group is significantly younger and different in gender/ethnic mix. In order to remove potential confounding effects, we included these variables in multivariable predictive models comparing controls and HD patients. Due to the speed of vaccine rollout we did not capture primary responses to vaccines as originally envisaged. As third vaccine doses were not routinely offered to healthy individuals at the time of study, we were unable to collect third vaccine data for controls. We did not perform virus neutralisation assays on the control group as others have already shown lower responses in patients receiving HD treatment, using the same assay [27].

In summary, we find that HD patients induce antigen-specific antibody responses that differ based on how the antigen is encountered. Our results suggest that, despite exhibiting a high infection risk suggesting immune dysfunction, patients requiring HD can mount effective recall immune responses, and that mRNA vaccine platforms may potentially enhance the functionality of antibody responses in this patient population. As such, we suggest that greater study of immune responses in this vulnerable patient population, particularly utilising the recent advances in mRNA vaccine technology, would allow development of strategies to optimise outcomes after vaccination. One area of immediate relevance where this could be tested include pathogens of clinical interest where current vaccination strategies yield poor or inconsistent results, for example seasonal influenza and hepatitis B [9].

## Methods

### Patient selection and data collection

Patients established on HD were recruited from two UK centres – University Hospital Birmingham Foundation Trust (UHBFT) and University Hospitals of Leicester NHS Trust (UHL). Patients from 12 UHBFT satellite dialysis units were recruited to a prospective observational study of immune responses to SARS-CoV-2 vaccination (Coronavirus Immunological Analysis (CIA) study, ethical approval granted by NorthWest-Preston Research Committee, ref 20/NW/0240). Control subjects were recruited from UHBFT and University of Birmingham employees (through internal advertising) and the general public, as part of the CIA study. UHL patients requiring dialysis were recruited to the “Phenotyping Seroconversion Following Vaccination Against COVID-19 In Patients On Haemodialysis” study (ethical approval granted by West Midlands-Solihull Research Ethics Committee: ref 21/WM/0031). Aspects of this study have been published previously [27, 53]. Only individuals over the age of 18 and eligible for SARS-CoV-2 vaccination were approached for participation in the study. Prisoners and individuals that did not have capacity as defined by the Mental Capacity Act were excluded. Written informed consent was obtained from all subjects involved in the study.

UHBFT HD patients underwent weekly screening for SARS-CoV-2 infection by PCR testing of nasopharyngeal swabs as part of standard clinical care. UHBFT HD patients that returned a positive PCR result were routinely dialysed in a central “COVID cohort” dialysis centre for 14 days, with regular blood tests and clinical reviews as standard of care. UHL HD patients were only tested if they developed symptoms compatible with SARS-CoV-2 infection. No prospective data collection for incidence of SARS-CoV-2 infection was performed in controls. Demographics, SARS-CoV-2 vaccination status, laboratory and clinical data were collected from electronic patient records. Immunosuppression was defined as current use of immunosuppressant medication (e.g. prednisolone >5mg per day or equivalent dose of another steroid, tacrolimus, mycophenolate, azathioprine), cyclophosphamide/methotrexate/plasma exchange in last 6 months or immunosuppressive monoclonal antibody in last 12 months. Previous SARS-CoV-2 exposure was defined as confirmed positive nasopharyngeal SARS-CoV-2 PCR prior to start of study and/or detection of anti-nucleocapsid (anti-N) antibody at study entry.

### Serological analysis

Serum samples were collected 21-28 days after the second SARS-CoV-2 vaccination in all subjects and 21-28 days after third vaccine dose only in HD patients. Sera were analysed for antibody directed against the SARS-CoV-2 Spike protein receptor-binding domain (S-RBD) and nucleocapsid (N) using an established automated electrochemiluminescence assay (Elecsys Anti-SARS-CoV-2 S and N, Roche diagnostics) [54]. Seropositivity was defined as anti-S levels greater than 0.8U/ml.

Binding of complement components to anti-S antibody was assessed using a solid phase C1q-binding assay and C4b/3b/5b complement deposition assay as described previously [26]. Briefly, 96 well microtiter plates were coated with 0.1ug/ml HexaPro Wuhan SARS-CoV-2 S protein isolated from transfected HEK293F cells [55]. After blocking, diluted heat inactivated test sera (56 degrees C, 30 mins) were added to the plate and incubated for 1hr at room temperature (RT). After washing, a standardised complement source (pooled SARS-CoV-2 negative normal human serum) was added to each well for 1hr at 37degrees. Plates were incubated with monoclonal antibodies directed against complement proteins C1q, C3b, C4b and C5b, and the signal amplified using HRP-conjugated secondary antibodies and/or the Perkin Elmer ELAST amplification kit as per manufacturer instructions. Plates were developed using TMB Core (Bio-Rad) and reaction stopped with H_2_SO_4_. Optical density (OD) was read at 450 nm using a SpectraMax ABS Plus plate reader. The pooled mean OD of negative control wells for each assay was set as the “detection threshold” (0.5 for C1q; 0.1 for C4b, C3b and C5b).

Anti-viral neutralising antibody (nAb) activity against SARS-CoV-2 were analysed in only HD patient sera collected 21-28 days after second and third vaccines using high throughput live-virus neutralisation assays as previously described [56]. Briefly, neutralisation of live virus by serial dilutions of sera were evaluated using a SARS-CoV-2 isolate with a spike identical to variants of interest. In this study, we examined reactivity against two SARS-CoV-2 strains: Wuhan and Delta (B.1.617.2) isolates. The quantifiable assay range is from dilutions of 1:40 to 1:2560. Some sera display neutralising activity, but with an IC50 (the dilution at which 50% of infection is prevented) below this range. We defined “detectable inhibition” as sera displaying inhibition at the 1:40 dilution or higher.

### Statistical analysis

Statistical analysis of data was performed using Prism v9 (GraphPad) and SPSS v26 (IBM). Two-sided tests were used throughout, with p values of 0.05 or less considered significant. Categorical variables were compared using Chi^2^ or Fisher’s exact tests. For continuous variables, unpaired comparisons were made using Mann-Whitney U test and Wilcoxon signed-rank test was used for paired comparisons. When comparing multiple groups, Kruskal-Wallis test was used with post-hoc Dunn’s multiple comparisons testing.

Correlation analysis was performed using Spearman rank test. Multivariable linear and logistic regression modelling were used to examine predictors of antibody quantity/quality and subsequent SARS-CoV-2 infection incidence. Non-parametric continuous variables were log_10_ transformed prior to inclusion in regression modelling.

## Acknowledgements

We would like to thank all of the dialysis and renal research nursing staff who were involved in recruitment and sample collection at both UHBFT and UHL sites. We would also like to thank James Hodson for his advice on statistical methods and multivariable modelling for this study, together with Drs Edward Carr and Rupert Beale for their expertise with SARS-CoV-2 neutralisation assays.

