## Supplemental data for "Different vaccine platforms result in distinct antibody responses to the same antigen in haemodialysis patients"

**Supplementary Table 1. Predictors of anti-S antibody levels in HD patients after 2 vaccine doses.** Univariable and multivariable linear regression analysis shown. Multivariable model included all parameters shown. Dependent variable (anti-S antibody levels) was $log_{10}$ transformed for Normality. P values <0.05 considered significant and highlighted in bold typeface.

| | **Univariable analysis** | | **Multivariable analysis** | |
|---|---|---|---|---|
| | Beta coefficient (95% CI) | p value | Beta coefficient (95% CI) | p value |
| Age (decades) | 1.00 (0.98, 1.02) | 0.84 | 1.00 (0.98, 1.03) | 0.64 |
| Female gender | 0.71 (1.35, 2.75) | 0.29 | 0.87 (0.50, 1.53) | 0.64 |
| White ethnicity | 0.33 (0.19, 0.62) | **<0.001** | 0.58 (0.32, 1.04) | 0.07 |
| Diabetes mellitus | 1.29 (0.66, 2.45) | 0.46 | 0.89 (0.50, 1.60) | 0.70 |
| Immunosuppression | 0.23 (0.08, 0.69) | **0.009** | 0.22 (0.09, 0.59) | **0.003** |
| Previous SARS-CoV-2 infection | 4.07 (2.19, 7.59) | **<0.001** | 4.61 (2.60, 8.17) | **<0.001** |
| mRNA vaccine | 5.24 (2.82, 9.77) | **<0.001** | 6.14 (3.48, 10.79) | **<0.001** |

**Supplementary Figure 1 – Serum virus neutralization activity in HD patients 21-28 days after 2 vaccine doses.**

**A:** Virus neutralisation activity of sera against Wuhan and Delta strains – data split between viral vector (grey symbols) and mRNA (blue symbols) vaccinees and groups split by previous SARS-CoV-2 infection (filled symbols denote previous infection, unfilled symbols – no infection). IC50 of 40 was used as threshold of activity (dashed green line). Percentages indicate proportions of individuals with detectable neutralization activity in each subgroup. **B:** Comparison of anti-S antibody levels between Delta nAb positive (filled symbols) and negative (unfilled symbols) individuals, split by previous infection status. Mann Whitney U test p values shown.

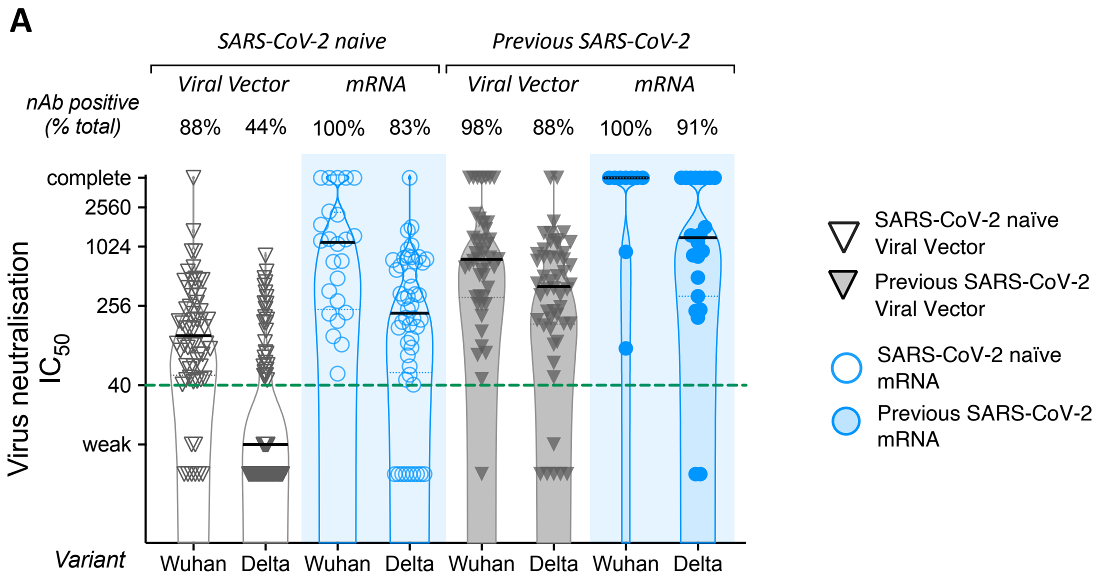


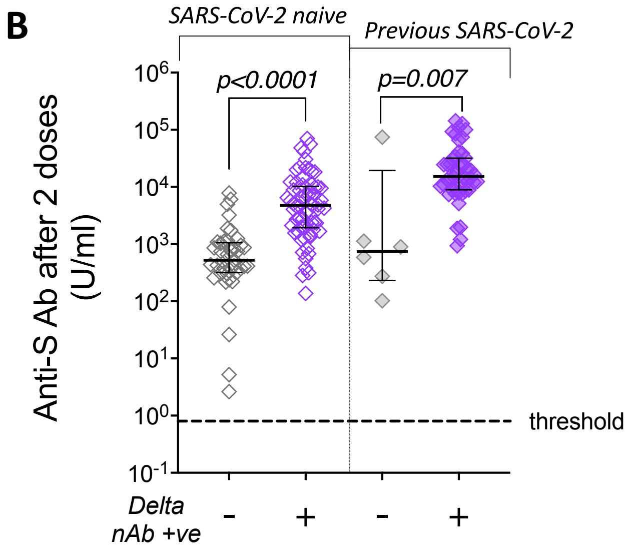

**Supplementary Table 2. Predictors of post-vaccination SARS-CoV-2 infection in HD patients after 2 vaccine doses.**

Univariable and multivariable logistic regression analysis shown. Dependent variable was incidence of SARS-CoV-2 infection after 2 doses of vaccine (before administration of third dose). Multivariable model included age, gender, ethnicity, diabetes, immunosuppression, HD centre and vaccine type (statistics shown for model with highly functional antibody – Delta nAb +ve with high complement binding). Previous SARS-CoV-2 exposure was significantly co-linear with all measures of antibody functionality and was, therefore, not included in the model.

*Measures of antibody functionality (Delta nAb, complement binding and combined nAb and complement binding) were added individually to the model as they demonstrated significant co-linearity. P values <0.05 considered significant and highlighted in bold typeface.

*Abbreviations: OR - Odds ratio; HD – haemodialysis, nAb – neutralizing antibody*.

| | **Univariable analysis** | | **Multivariable analysis** | |
|---|---|---|---|---|
| | OR (95% CI) | p value | OR (95% CI) | p value |
| Age (decades) | 1.07 (0.72, 1.59) | 0.73 | 1.13 (0.68, 1.87) | 0.65 |
| Female gender | 0.75 (0.27, 2.11) | 0.59 | 0.47 (0.14, 1.60) | 0.23 |
| White ethnicity | 0.90 (0.34, 2.40) | 0.84 | 0.37 (0.11, 1.29) | 0.12 |
| Diabetes mellitus | 0.70 (0.25, 1.96) | 0.50 | 0.36 (0.10, 1.28) | 0.11 |
| Immunosuppression | 0.56 (0.07, 4.48) | 0.58 | 0.28 (0.03, 2.89) | 0.29 |
| HD centre | 0.97 (0.36, 2.58) | 0.95 | 1.79 (0.29, 11.1) | 0.53 |
| mRNA vaccine | 1.06 (0.39, 2.88) | 0.91 | 1.11 (0.17, 7.04) | 0.92 |
| Detectable C1q-C5b binding* | 0.32 (0.11, 0.88) | **0.028** | 0.21 (0.06, 0.70) | **0.01** |
| Delta nAb +ve* | 0.27 (0.10, 0.77) | **0.014** | 0.15 (0.04, 0.52) | **0.003** |
| Delta nAb +ve with high complement binding* | 0.49 (0.27, 0.87) | **0.015** | 0.34 (0.17, 0.70) | **0.003** |
